# Human vaginal microbiota colonization is regulated by female sex hormones in a mouse model

**DOI:** 10.1101/2023.09.27.559827

**Authors:** Nuzhat Rahman, Firoz Mian, Charu Kaushic

**Author notes:** **Correspondence:** Charu Kaushic, McMaster Immunology Research Center, Department of Medicine, Michael G. DeGroote Center for Learning and Discovery, McMaster University, Hamilton, Ontario, Canada L8N 3Z5.

## Abstract

Clinically, a *Lactobacillus* rich vaginal microbiota (VMB) is considered optimal for reproductive outcomes, while a VMB populated by anaerobes is associated with dysbiosis and the clinical condition bacterial vaginosis (BV), which is linked to increased susceptibility to sexually transmitted infections and adverse reproductive outcomes. Mouse models that mimic eubiotic and dysbiotic VMB are currently lacking but could play a critical role in improving protective interventions. In this study, probiotic, eubiotic, and dysbiotic models were first developed in normal mice, using probiotic strains *Lactobacillus rhamnosus* GR-1 and *Lactobacillus reuteri* RC-14, eubiotic *Lactobacillus crispatus*, or dysbiotic *Gardnerella vaginalis* strains. Following a single administration, *L. rhamnosus* and *L. reuteri* persisted in the mouse vaginal tract for up to eight days, *L. crispatus* persisted for up to three days, and *G. vaginalis* persisted for up to two days, as measured by quantitative plating assays and qPCR. Colonization of *G. vaginalis* was facilitated by the presence of mucin. Endogenous sex hormones were manipulated by either ovariectomizing (OVX) mice or administering 17β-estradiol and progesterone pellets in OVX mice. The lack of endogenous hormones in OVX mice dramatically decreased VMB bacterial load compared to normal mice. None of the exogenous bacteria including *Lactobacilli* could colonize OVX mice for more than 24 hours. Treatment with 17β-estradiol but not progesterone restored the endogenous microbiota and colonization with *Lactobacilli* and *G. vaginalis*. Interestingly, 17β-estradiol treated mice had significantly increased levels of glycogen, compared to OVX and progesterone-treated mice. These results suggest there is a dynamic interaction between sex hormones and the VMB, which can affect bacterial diversity and the ability for a VMB to colonize.

## Introduction

The female reproductive tract (FRT) is colonized by an endogenous collection of microbes, termed the vaginal microbiota (VMB), which exists in a mutualistic relationship with the host (Chen, 2021). The VMB can be characterized into five different community state types (CST), with four of them being dominated by different *Lactobacillus* species whereas the fifth comprises of lower proportions of *Lactobacillus* and higher proportions of anaerobic organisms (Ravel et al., 2011). A eubiotic VMB is associated with CST I, which is characterized by low diversity and is *Lactobacillus crispatus* dominant (Ravel et al., 2011; Ghartey et al., 2014; Chee et al., 2020). A *L. crispatus* rich VMB has a low vaginal pH as a result of the large amount of lactic acid produced by the bacteria (Ravel et al., 2011). A low vaginal pH reduces pathogen viability and is associated with a protective outcome against pathogen exposure in the FRT (Miller et al., 2016). CST IV is considered to be a dysbiotic VMB and is composed of a diverse community structure. A dysbiotic VMB is comprised of strictly anaerobic bacteria such as *Gardnerella vaginalis*, a bacterial species that is commonly seen in patients with the clinical condition bacterial vaginosis (BV) (Morrill et al., 2020). *G. vaginalis* produces several virulence factors such as sialidase A and vaginolysin (Pleckaityte et al., 2012), and has been linked to biofilm formation (Patterson et al., 2010), allowing it to often outcompete *Lactobacillus* species in the vaginal tract. Crucially, women with BV are at an increased risk of acquiring sexually transmitted infections (STI) such as HIV-1 (Harold et al., 1999; Cherpes et al., 2003; Rebecca et al., 2010). The majority of data showing effects of the VMB on reproductive health comes from clinical studies and is correlative. There is a lack of VMB associated animal models in the literature that can be used for understanding the mechanism underlying the complex role the VMB plays in female reproductive health.

Many factors are known to affect VMB composition such as diet, ethnicity, antibiotic use, and importantly, sex hormones (Lehtoranta et al., 2022). A metanalysis examining the changes in the VMB throughout various phases of a women’s gynecological lifecycle found distinct microbial profiles at different stages of a woman’s reproductive life (Kaur, 2019). As the shifts between these gynecological stages are largely regulated by fluctuations in sex hormones, there is an underlying relationship between hormones and the changes in the VMB throughout a woman’s life. At puberty and pregnancy, when there is a major shift in endogenous hormones, there is an increase in glycogen deposition in the vaginal walls, enabling glycogen degrading *Lactobacillus* species to grow, which is considered optimal (Amabebe and Anumba, 2018). There is a shift from a low diversity to high diversity VMB when circulating sex hormones decrease such as during menopause (Oliveira et al., 2022), which is also implicated in increased susceptibility to other diseases such as heart disease and stroke, gynaecological malignancies, osteoporosis, and various genitourinary conditions (Elias and Sherris, 2003). Given the critical role of sex hormones on the VMB, it is important to take this into consideration when developing models.

A few studies have attempted to colonize mice with human VMB (Wolfarth et al., 2020), or BV-associated bacteria (Gilbert et al., 2019), however comprehensive studies that colonize mice with eubiotic and dysbiotic human VMB species have not yet been published. In this study, we successfully colonized normal female mice with human derived VMB species. A eubiotic model was developed with *L. crispatus* and a dysbiotic model was developed with *G. vaginalis*. Dysbiosis is associated with a heterogenous VMB, however since *G. vaginalis* is the most common bacteria seen in BV patients (Chen, 2021), this was the primary focus of the current study. As a positive control, a model with probiotic *Lactobacillus* species was also developed. To attempt to improve upon colonization, bacteria were supplemented with the nutrient sources glycogen or mucin. Given the critical role of hormones in VMB colonization, hormones were altered in mice to determine if there are relationships between sex hormones and VMB colonization *in vivo*. We looked at the VMB throughout the mouse estrus cycle, depleted all sex hormones by ovariectomizing (OVX) mice, and determined the effects of individual sex hormones by treating OVX mice with 17β-estradiol, the primary form of estrogen circulating in women during reproductive years (Ryan, 1982), or progesterone. Estrogen was found to promote VMB colonization in mice. 17β-estradiol was associated with increased glycogen in the vaginal tract, a common nutrient source used by bacteria (Hertzberger et al., 2022), which could explain the relationship between increased VMB colonization in 17β-estradiol treated mice. Collectively, our study successfully developed novel *in vivo* mouse models that harbour human-derived VMB species in hormone-unaltered and hormone altered mice. These models will serve as invaluable tools in studying the relationship between the VMB and female reproductive health.

## Materials and Methods

### Mice

Six–eight-week-old female C57BL/6 mice were obtained from Charles River Laboratories and housed in the Central Animal Facility at McMaster University. Mice were maintained under specific pathogen-free and standard temperature-controlled conditions that followed a 12h light/dark cycle and fed low-fat mouse chow. Mice were allowed one week after arrival to acclimate prior to experimental use. All mouse studies performed were approved by and were in compliance with the Animal Research Ethics Board at McMaster University.

### Bacteria Stock Preparation

*Lactobacillus crispatus* SJ-3C-US (PTA10138) from American Type Culture Collection (ATCC) was provided by Dr. Nuch Tanphaichitr (University of Ottawa, Canada). Probiotic *Lactobacillus* species *L. reuteri* (RC-14) and *L. rhamnosus* (GR-1) were received in the form of stab-cultures from the laboratory of Dr. Gregor Reid (Western University, Canada). *Gardnerella vaginalis* ATCC 14019 was purchased from ATCC. *L. rhamnosus, L. reuteri*, and *L. crispatus* were grown in ATCC medium 416 (Lactobacillus MRS broth/agar) in anaerobic conditions using the GasPak™ EZ Anaerobe Container System (Becton, Dickinson and Company, USA, Cat #260001) at 37°C. *G. vaginalis* was grown in ATCC medium 1685 (NYC III medium) in anaerobic conditions using the GasPak™ EZ Anaerobe Container System at 37°C. Stocks suspended in 20% glycerol were prepared for each bacterium and stored at -80°C for future use. To determine the bacterial stock concentration, serially diluted stocks were plated onto MRS agar plates (*L. rhamnosus, L. reuteri*, and *L crispatus*) or tryptic soy agar supplemented with 5% sheep’s blood *(G. vaginalis*) to determine Colony Forming Units (CFU)/mL by the Miles and Misra technique (Miles et al., 1938).

### Collection of Vaginal Washes

Two 30 μL volumes of sterile PBS were pipetted in and out of the mouse vagina 5-7 times, resulting in a total of 60 μL being collected. If the samples were used to check to bacterial colonization in the mice or to stage the mice in their estrus cycle, they were used right away. If they were collected to isolate DNA, then the samples were stored at -80°C.

### Colonization of Mice

Bacteria were grown in their respective media for 24 h. After 24 h of growth, the bacteria were spun down at 4000 rpm for two minutes, washed with PBS, and spun down once again. After the wash, bacteria were resuspended in PBS at 10^7^ CFU for one mouse in 25 μL volumes. Depending on the number of mice being colonized, a greater volume was prepared. To colonize the mice, 25 μL of the bacteria was pipetted into the vaginal canal. The mice were then held facedown to allow for the bacteria to persist in the vaginal canal for at least one minute. To mimic a eubiotic human VMB, mice were inoculated with *L. crispatus* at a concentration of 10^7^ CFU. A probiotic model was also developed by inoculating *L. rhamnosus* GR-1 and *L. reuteri* RC-14 together in equal concentrations of 5 × 10^6^ CFU to give a final concentration of 10^7^ CFU. *Lactobacillus* species were supplemented with 5 μL of 20mg/mL glycogen (0.1 mg) as a nutrient source in some experiments. To create the dysbiotic model, mice were inoculated with *G. vaginalis* at 10^7^ CFU. *G. vaginalis* was supplemented with 5 μL 10 mg/mL (0.05 mg) mucin in some experiments. To evaluate colonization, vaginal washes were collected from the mice and quantitative plating assays were performed as described below.

### DNA Extraction from Vaginal Washes

Vaginal washes were collected from mice and frozen at -80°C for DNA extraction. DNA was isolated from vaginal washes or from cultured bacteria using the DNeasy™ Blood & Tissue Kit (Qiagen, Netherland, Cat #69506). DNA was isolated as per the manufacturer’s instructions, including a primary digestion step using lysozyme to target the gram positive bacteria cell wall (Gill et al., 2016; Martzy et al., 2019), the primary type of bacteria in the mouse VMB (Vrbanac et al., 2018). For the latter method, an enzymatic lysis buffer consisting of 20 mM Tris·Cl (pH 8.0), 2 mM sodium EDTA, and 1.2% Triton® X-100 was prepared, and immediately before use, 20 mg/ml of lysozyme was added. The wash or cultured bacteria was pelleted and resuspended in 180 μl enzymatic lysis buffer and incubated for 30 minutes at 37°C. Following enzymatic lysis, DNA extraction was completed as per the manufacturer’s instructions for the DNeasy™ Blood & Tissue Kit.

### Quantitative PCR (qPCR)

qPCR of 16S rRNA gene was performed to assess overall bacteria load (Rezki et al., 2016), as well as species DNA present in a sample using species specific primers for *L. crispatus* (Zozaya-Hinchliffe et al., 2010), *L. rhamnosus* (Kim et al., 2020), *L. reuteri* (Kim et al., 2020), and *G. vaginalis* (Zozaya-Hinchliffe et al., 2010) (Table 1). A master mix containing 12.5 μL RT^2^ SYBR^®^ Green qPCR master mix (Qiagen, Netherlands, Cat #330503), 0.25 μL forward primer (100 μM), 0.25 μL reverse primer (100 μM), and 7 μL water per well was prepared. 20 μL of the master mix was aliquoted to each well of a 96-well plate and 5 μL of template DNA was added accordingly. The annealing temperature was input as T_m_ – 5°C, where the lower of the two T_m_ from the forward and reverse primers was used. The reaction was run using the StepOne Plus™ Real-Time PCR System (ThermoFisher™, USA). Samples were run in triplicate and bacterial load or species load was assessed by analyzing the number of cycles required for the fluorescent signal to cross the threshold (ct value). This value was inversed in our figures to show a positive relationship for ease of readability.

**Table 1:**
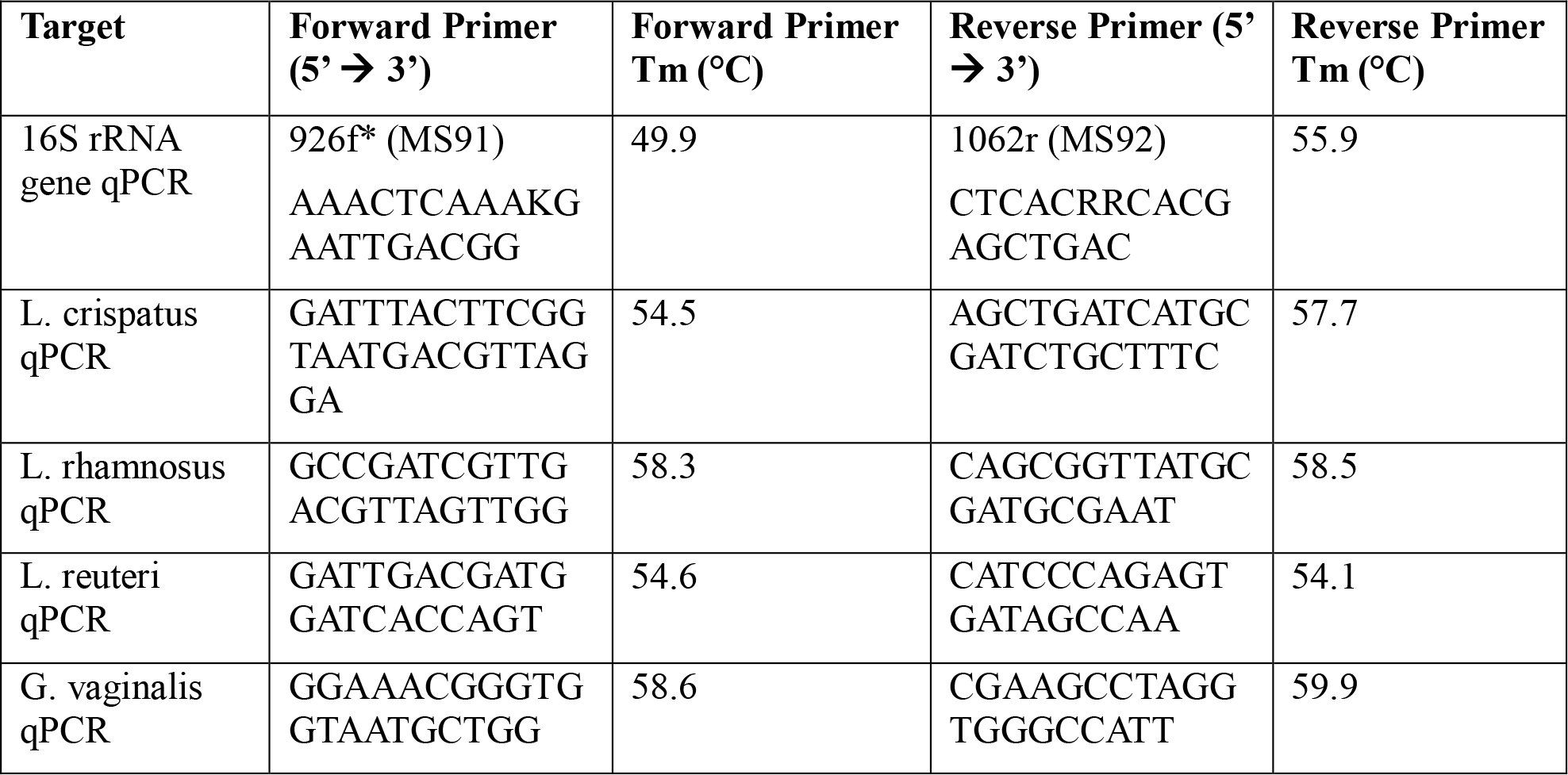
qPCR primers for the 16S rRNA gene, *Lactobacillus* species, and *G. vaginalis*.

### Quantification of VMB by Culture

Vaginal washes were collected from individual mice and serially diluted ten-fold from 10^−1^ to10^−6^. If *Lactobacillus* was the bacteria being screened, MRS plates were used and if *G. vaginalis* or endogenous bacteria were being screened, tryptic soy agar plates supplemented with 5% sheep’s blood were used. Plates were divided into 6 sections, and 10 μL of each dilution (sometimes including the undiluted sample) was pipetted dropwise onto the plate in duplicate. The plates were incubated in anaerobic conditions using the GasPak™ EZ Anaerobe Container System at 37°C. After 24 h, the number of colonies were counted using the dilution that isolated colonies could be identified and CFU/mL was calculated. Endogenous and exogenous colonies were identified by visually comparing colonies based on morphology from plating vaginal washes from untreated mice or from streaked bacterial stocks on plates, respectively.

### Ovariectomy (OVX)

Ovariectomies were performed to eliminate the effect of endogenous sex hormones in mice. Mice received the analgesic carprofen (5 mg/kg) subcutaneously and after 30 min, they were administered an intraperitoneal injection of ketamine (100-150 mg/kg) and xylazine (10 mg/kg). After the mice reached surgical plane, the surgical area was shaved, the mice received an intradermal injection of bupivacaine (4 mg/kg per incision site), and the surgical site was sterilized with iodine scrub and isopropyl alcohol. The ovaries were then removed through a small incision near the hind limbs and incision sites were sutured and stapled. 1 mL of saline was administered, and the mice recovered on a heat pad and were monitored until they were awake and able to move on their own. Mice received carprofen (5 mg/kg) for two days after surgery and were monitored for 5 days post surgery. Staples were removed 7-10 days post surgery and mice were allowed at least 1 week to recover before use.

### Hormone Treatments

Mice received the analgesic carprofen (5 mg/kg) subcutaneously and after 30 minutes, they were anaesthetized with isoflurane gas anesthesia. After the mice reached surgical plane, the surgical area was shaved, the mice received an intradermal injection of bupivacaine (4 mg/kg), and the surgical site was sterilized with iodine scrub and isopropyl alcohol. 10 mg progesterone 21-day release pellets (Innovative Research of America, USA, Cat #P-131-10MG-25) and 0.01 mg 17β-estradiol 21-day release pellets (Innovative Research of America, USA, Cat #E-121-0.01MG-25) were surgically inserted into the scruff of mice. These doses correspond to hormone levels measured during the estrous cycle (Bhavanam et al., 2008; Bagri et al., 2020) Mice received carprofen (5 mg/kg) for 2 days after surgery and were monitored for 5 days post surgery. Staples were removed 7-10 days post surgery and mice were allowed at least 1 week to recover before use.

### Immunohistochemistry of Vaginal Tissue

Mouse vaginal tissue was collected, placed in cassettes, and fixed in methacarn (60% methanol, 30% chloroform and 10% glacial acetic acid) for 72 h. Cassettes were transferred to 70% ethanol and samples were taken to McMaster Immunology Research Center (MIRC) Histology Core Facility for processing. The tissue was embedded, and slides were cut and mounted on microscope slides and stained with Mucin-1 (MUC-1) antibody (Abcam, United Kingdom, Cat #ab15481). Slides were scanned using the Leica Aperio Scanscope XT and viewed using Aperio ImageScope software.

### Statistical Analysis

All statistical analysis was done using GraphPad Prism version 9.4.1 (GraphPad Software, San Diego, CA). Two-way ANOVA with Tukey’s multiple comparisons was used to determine all statistical significance for quantitative plating assays. A one-way ANOVA with Tukey’s multiple comparisons or t-test was used in all qPCR experiments and ELISA.

## Results

### Normal female mice can be colonized by human VMB species

In order to develop a mouse model of human female microbiota, we first assessed if the vaginal tract of normal mice can be colonized by human VMB strains. We developed a eubiotic model with the physiologically relevant CST I bacteria *L. crispatus* and a dysbiotic model with the primary BV-associated bacteria *G. vaginalis*. A model using probiotic species *L. rhamnosus* GR-1 and *L. reuteri* RC-14 was also developed as a positive control, as probiotics are more robust and more likely to survive in non-optimal environments (Corcoran et al., 2005). For a general term usage, we refer to these bacterial species as exogenous bacteria, whereas the normal vaginal bacteria of the mice are referred to as endogenous. Female mice were inoculated intravaginally once with a total of 10^7^ CFU of RC-14 and GR-1 in equal concentrations, *L. crispatus* or *G. vaginalis*. PBS was administered as a negative control involving no exogenous bacteria. These groups of mice were then tracked for colonization using quantitative plating assays. As expected, no exogenous bacteria were detected in the PBS inoculated mice (Figure 1A). 6/6 (100%) mice in the probiotics group (Figure 1B), 5/6 (83%) mice in the *L. crispatus* group (Figure 1C), and 4/9 (44%) mice in the *G. vaginalis* group (Figure 1D), were successfully colonized for at least 24 h. Mice inoculated with probiotic species were colonized for a minimum of 2 days and a maximum of 8 days, with an average duration of 4.2 days. Mice given *L. crispatus* were colonized for two days minimum and three days maximum, with an average duration of colonization of 2.6 days. *G. vaginalis* treated mice were colonized for one day minimum and two days maximum, with an average duration of colonization of 1.75 days.

**Figure 1.**
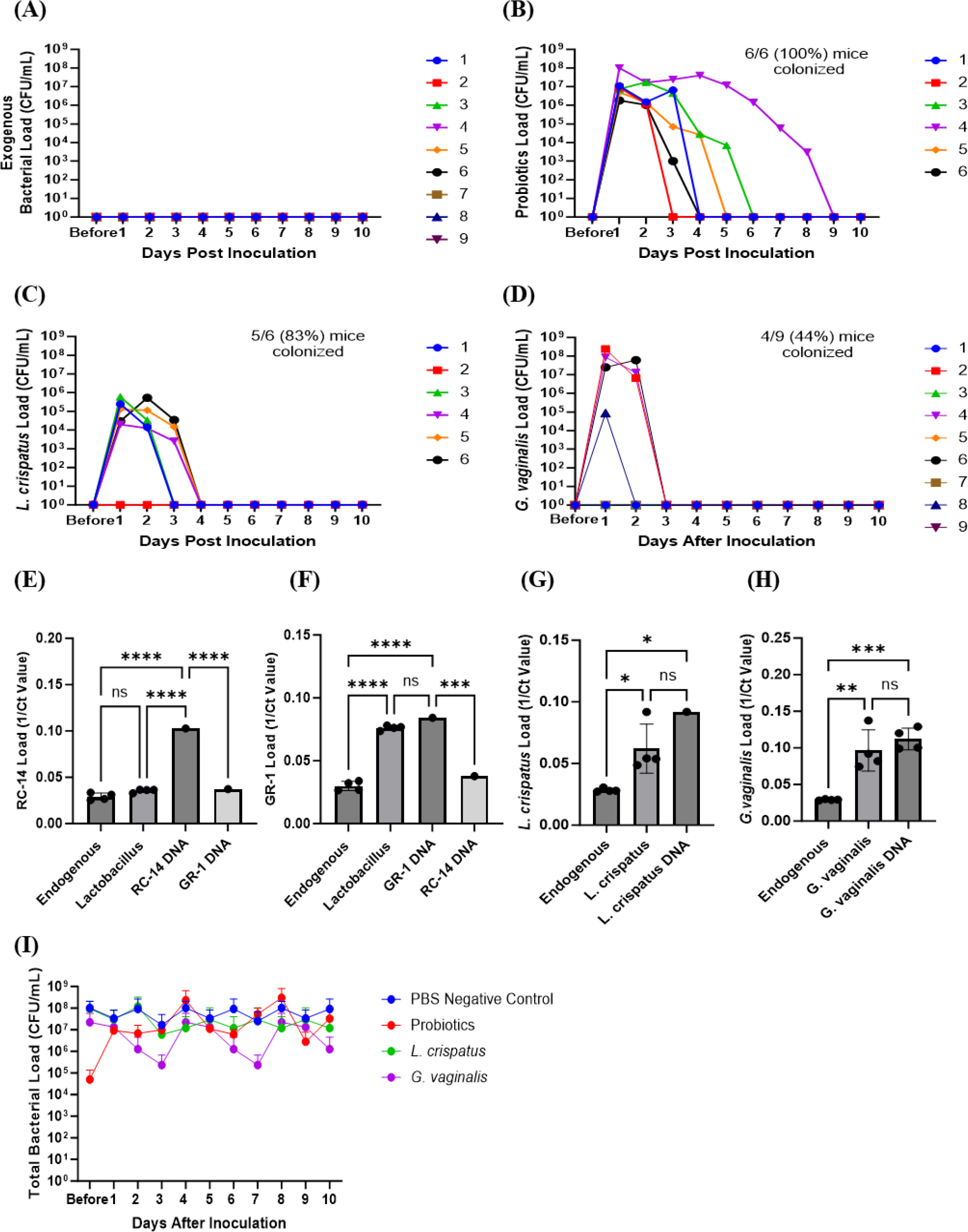
Normal mice colonized with *Lactobacillus* probiotic species, *L. crispatus*, and G. vaginalis. Female mice were inoculated once with a total of 10^7^ CFU *L. reuteri* RC-14 and *L. rhamnosus* GR-1 in equal concentrations, *L. crispatus, G. vaginalis*, or PBS as a negative control. Data are from n = 6-9 per group from one experiment representative of 3 independent experiments with similar results. Vaginal washes were collected up to 10 days post-inoculation, and bacteria colonies from the inoculated species types were counted for samples from PBS inoculated (A), probiotics inoculated (B), *L. crispatus* inoculated (C), and *G. vaginalis* inoculated (D) groups on agar plates. Different mice are denoted by different colored points. The data was analyzed using a two-way ANOVA with Tukey’s multiple comparisons, but no significance was found. From these plating assays, probiotic-looking colonies (n = 4 colonies), *L. crispatus*-looking colonies (n = 4 colonies), *G. vaginalis*-looking colonies (n = 4 colonies), and endogenous colonies (n = 4 colonies) were picked from the culture plates and DNA was isolated. Using this DNA, qPCR was performed with RC-14 specific qPCR primers (E), GR-1 specific qPCR primers (F), *L. crispatus* specific qPCR primers (G), and *G. vaginalis* specific qPCR primers (H), to determine what type of bacterial DNA was present. RC-14, GR-1, *L. crispatus*, and *G. vaginalis* stock DNA were used as positive controls. Data was analyzed using a one-way ANOVA (****p<0.0001, ***p<0.001, **p<0.01, *p<0.05). The total vaginal bacterial load was plotted from all groups in panel I. The data was analyzed using a two-way ANOVA with Tukey’s multiple comparisons, but no significant difference was found.

To validate that the bacteria species being counted on the plates were indeed *L. reuteri* or *L. rhamnosus, L. crispatus*, or *G. vaginalis* and not endogenous mouse vaginal species, qPCR was used for validation. DNA was isolated from colonies of endogenous species, as well as probiotics, *L. crispatus* or *G. vaginalis* colonies plated from vaginal washes and distinguished by colony morphology. Colonies identified as endogenous bacteria by plating did not show significant bacterial count in qPCR using probiotic specific primers (Figures 1E and 1F), *L. crispatus* specific primers (Figure 1G), or *G. vaginalis* specific primers (Figure 1H), indicating that the primers were specific, and colonies identified and counted as endogenous did not contain exogenous bacteria. For the probiotics inoculated mice, qPCR using RC-14 and GR-1 specific qPCR primers was performed. There was a significant difference between the Ct values for *Lactobacillus* colonies and RC-14 stock DNA when using RC-14 specific qPCR primers (Figure 1E), but no significant difference between the Ct values for *Lactobacillus* colonies and GR-1 stock DNA when using GR-1 specific primers, indicating that the colonies being counted were predominantly *Lactobacillus rhamnosus* GR-1. Thus, among the probiotic mixture, GR-1 was the predominant bacteria colonizing the probiotic inoculated mice (Figure 1F). Likewise, there was no significant difference between Ct values for *L. crispatus* or *G. vaginalis* colonies and *L. crispatus* (Figure 1G) or *G. vaginalis* (Figure 1H) stock DNA, indicating the colonies counted as *L. crispatus* or *G. vaginalis* in the corresponding bacteria inoculated mice were indeed those bacteria.

The total bacterial load including endogenous and exogenous colonies in all groups was compared using quantitative plating assays by counting colonies of both endogenous and exogenous species. No significant increase in total bacteria was observed post inoculation, indicating that the exogenous bacteria were displacing the endogenous bacteria (Figure 1I). Total bacterial load remained consistently ∼ 10^7^-10^8^ CFU/mL in all groups, suggesting that there is a finite niche for bacteria in the vaginal tract of mice.

### Exogenous mucin facilitated initial G. vaginalis colonization

Since not all mice were colonized equally with different bacteria, we wanted to test methods to improve colonization. Nutrient availability is an important determinant of what species colonize the VMB (Hood-Pishchany and Rakoff-Nahoum, 2021). Glycogen is a common nutrient source used by *Lactobacillus* species (Mirmonsef et al., 2014) and mucin is a common nutrient source used by *G*. vaginalis (Dupont, 2020; Vagios and Mitchell, 2021). Therefore, we tested if addition of mucin or glycogen at the time of inoculation could enhance colonization. Supplementation with mucin or glycogen in probiotic (*L. reuteri* or *L. rhamnosus*) treated or *L. crispatus* treated mice did not significantly facilitate or hinder colonization (Figures 2A-2D). 6/6 (100%) mice in the probiotics plus glycogen group and 5/6 (83%) of mice in the probiotics plus mucin group were successfully colonized (Figures 2A and 2B), which is a similar rate to colonization rates in no nutrient conditions (Figure 1B). The average duration of colonization in successfully colonized mice in the probiotics plus glycogen and mucin groups was 4 days and 2.2 days respectfully, indicating mucin might be decreasing the duration of colonization in mice. 5/6 (83%) mice in the *L. crispatus* plus glycogen group and 8/9 (89%) of the mice in the *L. crispatus* plus mucin group were colonized, with an average duration of colonization of 2.2 days and 2.3 days respectively (Figures 2C and 2D). These values are similar to no nutrient supplementation (Figure 1C), indicating nutrient supplementation did not help or hinder *L. crispatus* colonization. Glycogen supplementation did not enhance *G. vaginalis* colonization, with 2/9 (22%) of mice being initially colonized (Figure 2E). However, supplementation with mucin aided in initial *G. vaginalis* colonization, with 8/9 (89%) of mice being colonized for at least 24 hours (Figure 2F). This is double the number of mice compared to no mucin supplementation (Figure 1D), suggesting exogenous mucin administration could assist in *G. vaginalis* colonization in mice.

**Figure 2.**
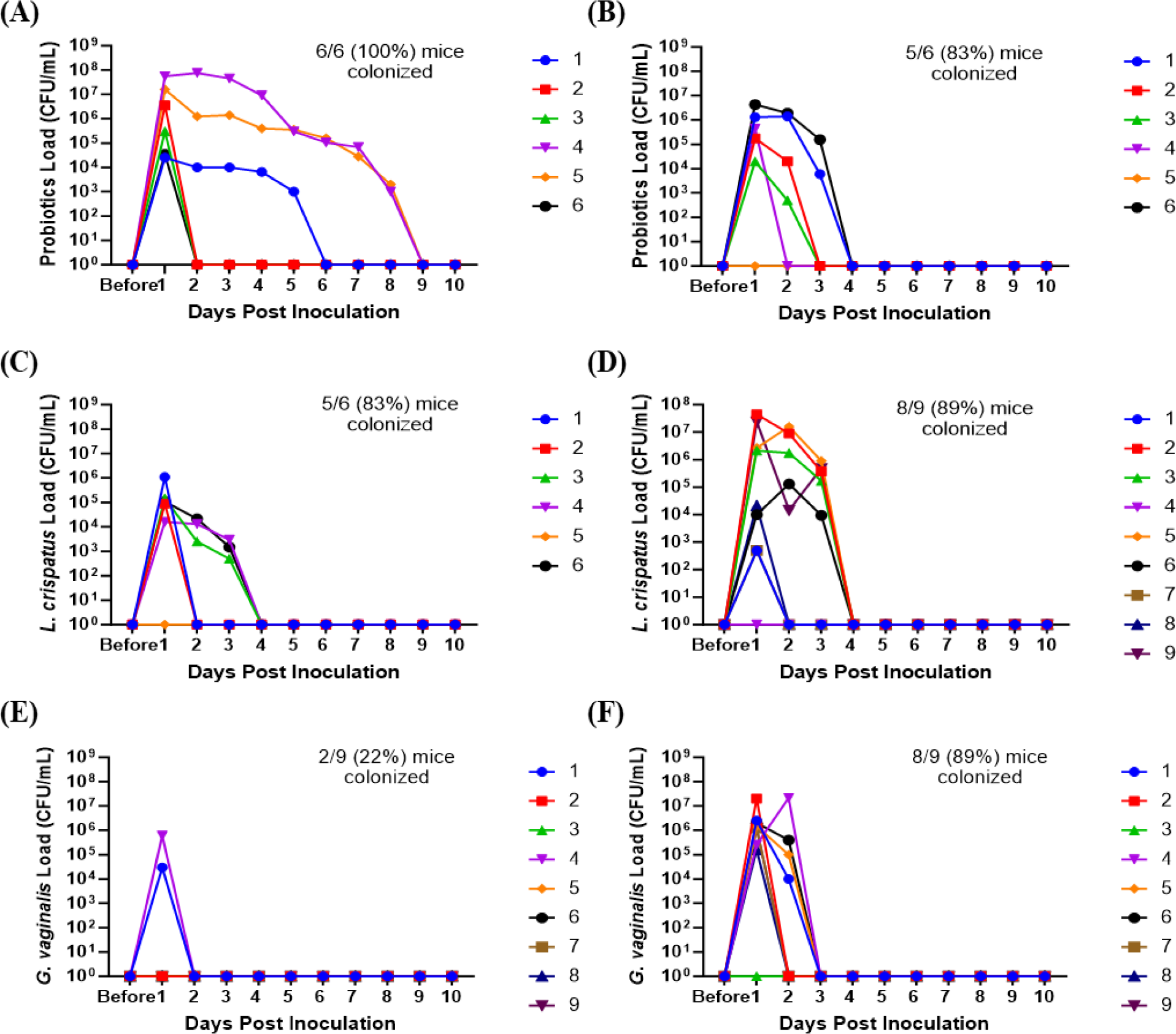
Mucin enhanced *G. vaginalis* colonization. Female mice were inoculated once with 10^7^ CFU of equal parts *Lactobacillus* probiotic species *L. reuteri* RC-14 and *L. rhamnosus* GR-1 plus glycogen (A), probiotic species plus mucin (B), *L. crispatus* plus glycogen (C), *L. crispatus* plus mucin (D), *G. vaginalis* plus glycogen (E), or *G. vaginalis* plus mucin (F). Vaginal washes were collected up to 10 days post-inoculation and assessed by quantitative plating assays. Specific bacterial colonies derived from mice inoculated with different bacterial species with either glycogen or mucin were counted on agar plates. Data indicate n = 6-9 per group, representative of 3 independent experiments. Different mice are denoted by different colored points. The data was analyzed using two-way ANOVA with Tukey’s multiple comparisons, but no significance was found.

### Mice were more likely to be colonized with exogenous bacteria in the estrus stage

Because significant differences were seen in successful initial colonization and duration of colonization between mice within the same treatment group, we considered other factors that could affect the colonization. The transition of women through menopause is marked by a gradual depletion of *Lactobacillus* species and an increase in anaerobic bacteria (Oliveira et al., 2022). Others have reported that mice given 17β-estradiol are more likely to be colonized by BV associated bacteria (Gilbert et al., 2019). Because of the identified relationships between hormones and the VMB, we first looked at the effect of the mouse estrus cycle on human VMB species colonization. During the estrus stage (the estrogen high phase of the reproductive cycle), all mice in probiotics treated, *L. crispatus* treated, and *G. vaginalis* treated were colonized (Figures 3A, 3D and 3G). In the progesterone high diestrus phase, 4/6 (67%) of probiotics treated mice (Figure 3B), 3/6 (50%) of *L. crispatus* treated mice (Figure 3E), and 1/6 (17%) of *G. vaginalis* treated mice (Figure 3H) were colonized, indicating that overall, estrogen may be facilitating colonization. When looking at the average of all mice in estrus or diestrus, there were no significant differences in probiotics (Figure 3C) and *L. crispatus* (Figure 3F) inoculated mice, however there was a significant difference one day post-inoculation in the *G. vaginalis* group (Figure 3I). This is most probably due to the stark difference in initial colonization success between estrus and diestrus mice in the *G. vaginalis* group.

**Figure 3.**
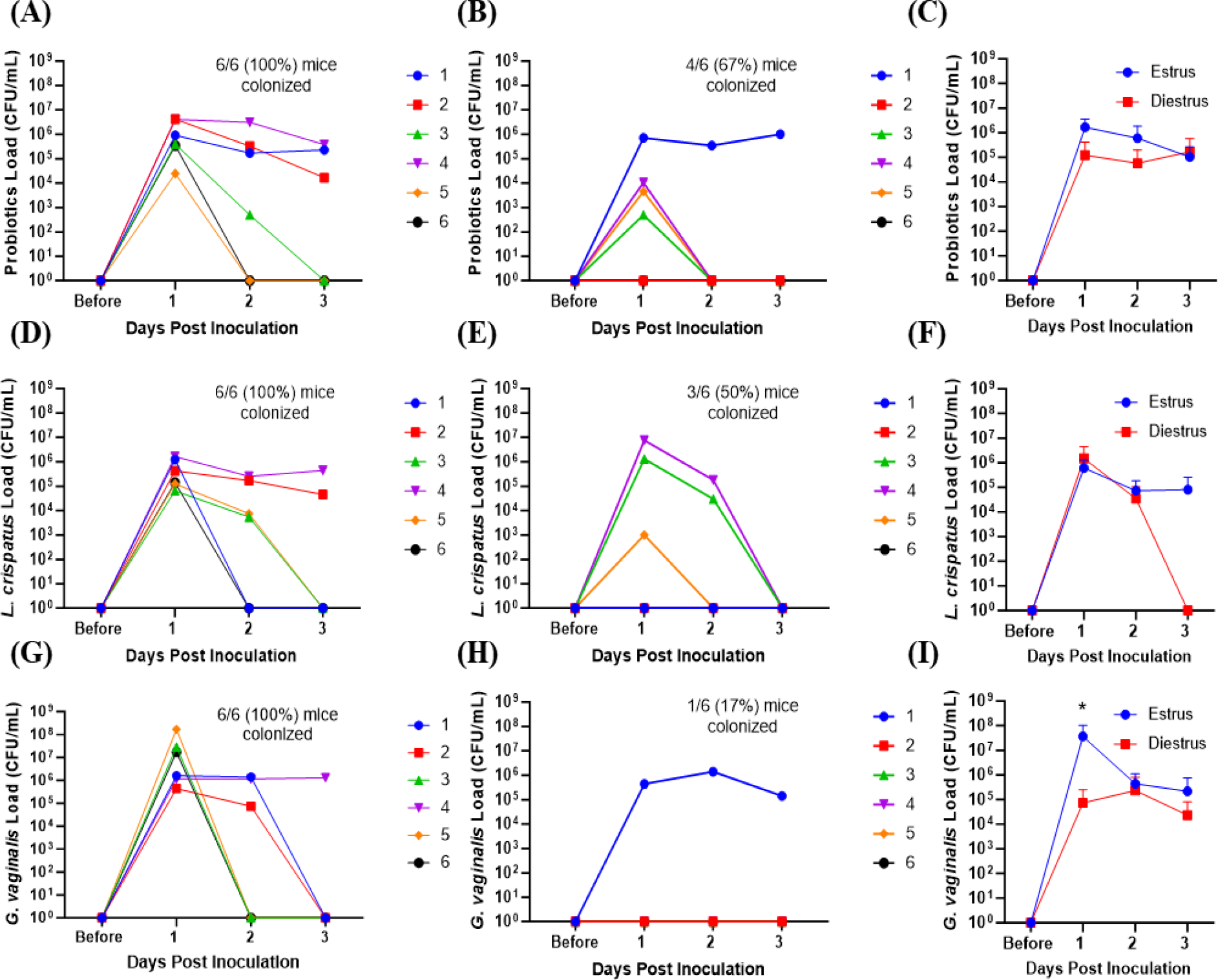
Mice were more likely to be successfully colonized by probiotic bacteria, *L. crispatus*, or *G. vaginalis* when they were in the estrus stage of the reproductive cycle. Probiotics load, *L. crispatus* load, or *G. vaginalis* load was assessed by quantitative plating assays before and after one vaginal administration of 10^7^ CFU of *Lactobacillus* probiotics species RC-14 and GR-1 in equal proportions, *L. crispatus*, or *G. vaginalis* in mice in estrus or diestrus on the day of inoculation (n=6 per group, representative of 2 independent experiments). Mice were monitored up to 3 days after inoculation and probiotics colonies in estrus (A), diestrus (B), and combined (C), *L. crispatus* colonies in estrus (D), diestrus (E), and combined (F), and *G. vaginalis* colonies in estrus (G), diestrus (H), and combined (I), were plotted. In panels A, B, D, E, G, and H, different mice are denoted by different colored points. The data in panels C, F, and I were analyzed using a two-way ANOVA with Tukey’s multiple comparisons (*p< 0.05).

### Hormone-depleted mice had decreased total bacterial load and were not colonized by exogenous human VMB species

We next examined the effect of eliminating endogenous sex hormones on the VMB of mice. Ovariectomies were performed on normal mice to eliminate the effect of endogenous sex hormones. Vaginal washes were collected from mice before and 1, 2 and 3 weeks after ovariectomies and quantitative plating assays were performed. There was a dramatic and significant decrease in the endogenous bacterial load of the mice after ovariectomy (Figure 4A). DNA was also isolated from vaginal washes collected from the same OVX mice before and 3 weeks post-OVX. qPCR using 16S rRNA gene specific primers to target total bacterial DNA was performed as a more quantitative measure of the amount of bacteria present before and after OVX. Vaginal washes showed a significantly decreased amount of bacterial DNA after ovariectomy compared to before (Figure 4B), validating the results from the plating assays.

**Figure 4.**
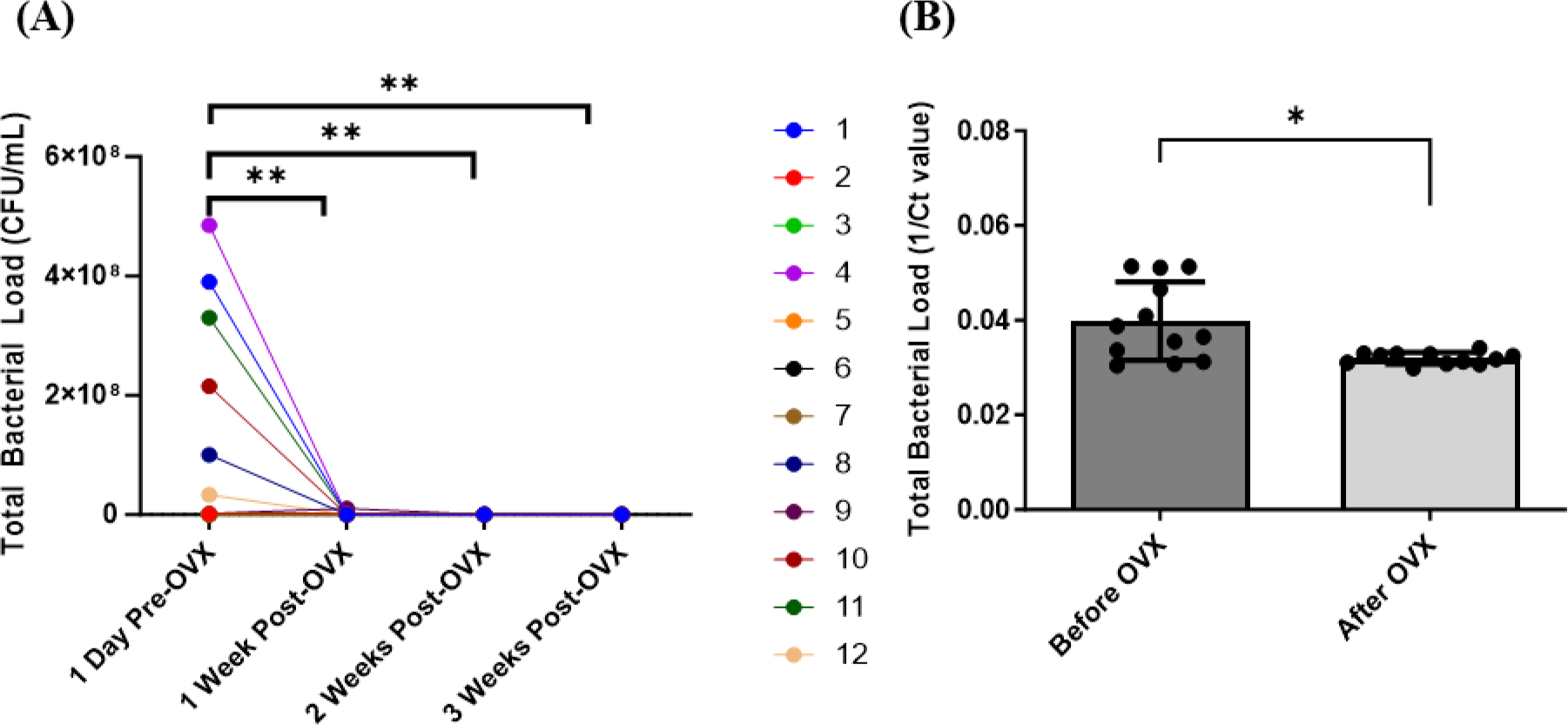
Mice showed decreased bacterial load post ovariectomies, as calculated by quantitative plating assays and qPCR. Vaginal washes were collected from individual mice 1 day before they were ovariectomized, as well as 1 week, 2 weeks, and 3 weeks post-OVX (n=12, representative of 2 independent experiments). Total bacterial load was assessed via quantitative plating assays (A) or 16S rRNA-specific qPCR (B). In panel A, different mice are denoted by different colored points. Data was analyzed using a two-way ANOVA with Tukey’s multiple comparisons (**p<0.01) or t-test (*p<0.05).

We also wanted to determine the ability for human VMB species to colonize and persist in hormone-depleted mice. A single inoculation, as well as 3 consecutive daily doses of probiotic *Lactobacillus* species RC-14 and GR-1, *L. crispatus*, or *G. vaginalis* was delivered to OVX mice at 10^7^ CFU. Probiotics, *L. crispatus*, and *G. vaginalis* colonies were counted for two days after the final inoculation. In the probiotics inoculated mice, there was brief colonization one day post inoculation, and then a rapid decrease by two days post inoculation in both single and triple inoculated mice, indicating that probiotic bacteria were not able to colonize OVX mice for any significant length of time (Figures 5A and B). On the individual mouse level, 5/6 (83%) of the mice in the single inoculated group (Figure 5A) and 8/8 (100%) of mice in the triple inoculated group (Figure 5B) were colonized. The total bacteria count only reached a maximum concentration of ∼10^3^ CFU/mL, which is 100,000 times lower than the ∼10^8^ CFU/mL reached in normal mice (Figure 1I). Similar to probiotics inoculated mice, the *L. crispatus* group had brief colonization one day post-inoculation at ∼10^3^ CFU/mL, and no bacteria was detected two days post inoculation in single and triple inoculated mice (Figures 5C and 5D). On the individual mouse level, 4/6 (83%) of the mice in the single (Figure 5C) and triple inoculated (Figure 5D) groups were colonized. In the *G. vaginalis* inoculated mice, no mice were colonized in the single inoculated group (Figure 5E) and bacteria was detected in 2 mice in the triple inoculated group, but once again, only for one day (Figure 5F). Overall, these results suggest the absence of hormones makes the vaginal environment inhospitable to both endogenous and exogenous bacteria, which could explain why the human VMB species we administered were not able to colonize, even temporarily.

**Figure 5.**
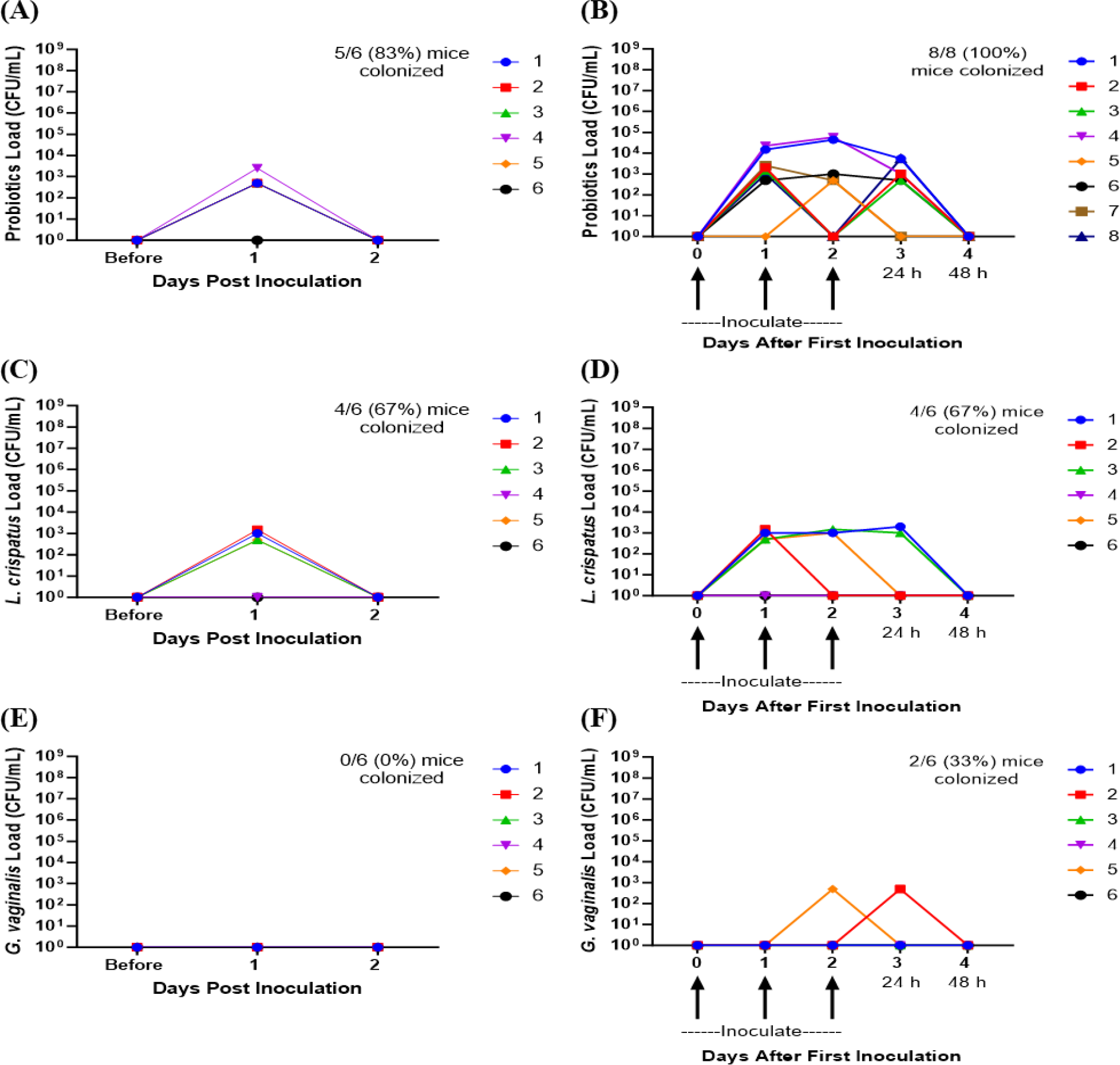
Ovariectomized mice did not get colonized with probiotic *Lactobacillus* species, *L. crispatus*, or *G. vaginalis* for more than 24 h after one inoculation and multiple inoculations. Female mice were ovariectomized and allowed to recover for 10 days. Mice were then inoculated once or three times consecutively every 24 h with a total of 10^7^ CFU *L. reuteri* RC-14 and *L. rhamnosus* GR-1 in equal concentrations, *L. crispatus*, or *G. vaginalis* (n=6 per group, representative of 2 independent experiments). Vaginal washes were collected up to 2 days post most recent inoculation, and probiotics load, *L. crispatus* load, or *G. vaginalis* load were assessed by quantitative plating assays. Probiotics colonies in mice inoculated once (A) or three times (B), *L. crispatus* colonies in mice inoculated once (C) or three times (D), or *G. vaginalis* colonies in mice inoculated once (E) or three times (F) were counted and plotted. Different mice are denoted by different colored points. In panels B, D, and F, 24 h or 48 h denote the time passed after the most recent inoculation. The data was analyzed using two-way ANOVA with Tukey’s multiple comparisons, but no significance was found.

### Hormone-depleted mice treated with estrogen had significantly increased bacterial load and were colonized by human VMB species

We next examined the effects of 17β-estradiol or progesterone given to OVX mice to assess the effect of these hormones on the VMB in mice. Ten days after mice recovered from ovariectomies, either a 10 mg 21-day release progesterone pellet or a 0.01 mg 17β-estradiol 21-day release pellet was inserted subcutaneously into the mice to recapitulate hormone levels during the estrus cycle (Bhavanam et al., 2008; Bagri et al., 2020). Ten days after the hormone pellet insertion, mice were inoculated once with a total of 10^7^ CFU probiotics RC-14 and GR-1, *L. crispatus, G. vaginalis*, or PBS as a no exogenous bacteria negative control. The bacterial load of the mice before hormone treatments (OVX mice) was very low (Figures 6A and 6B), similar to what was seen in previous experiments (Figures 4 and 5). After treatment with progesterone, there was a slight increase in bacterial load compared to OVX mice, but the difference was not statistically significant (Figure 6A). After treatment with 17β-estradiol, there was a statistically significant increase in bacterial load compared to OVX mice, indicating that 17β-estradiol may play a role in human VMB establishing in mice (Figure 6B). Of note, 17β-estradiol treated mice had a bacterial load in the ∼10^7^-10^8^ CFU/mL range (Figure 6B), which is similar to normal mice (Figure 1).

**Figure 6.**
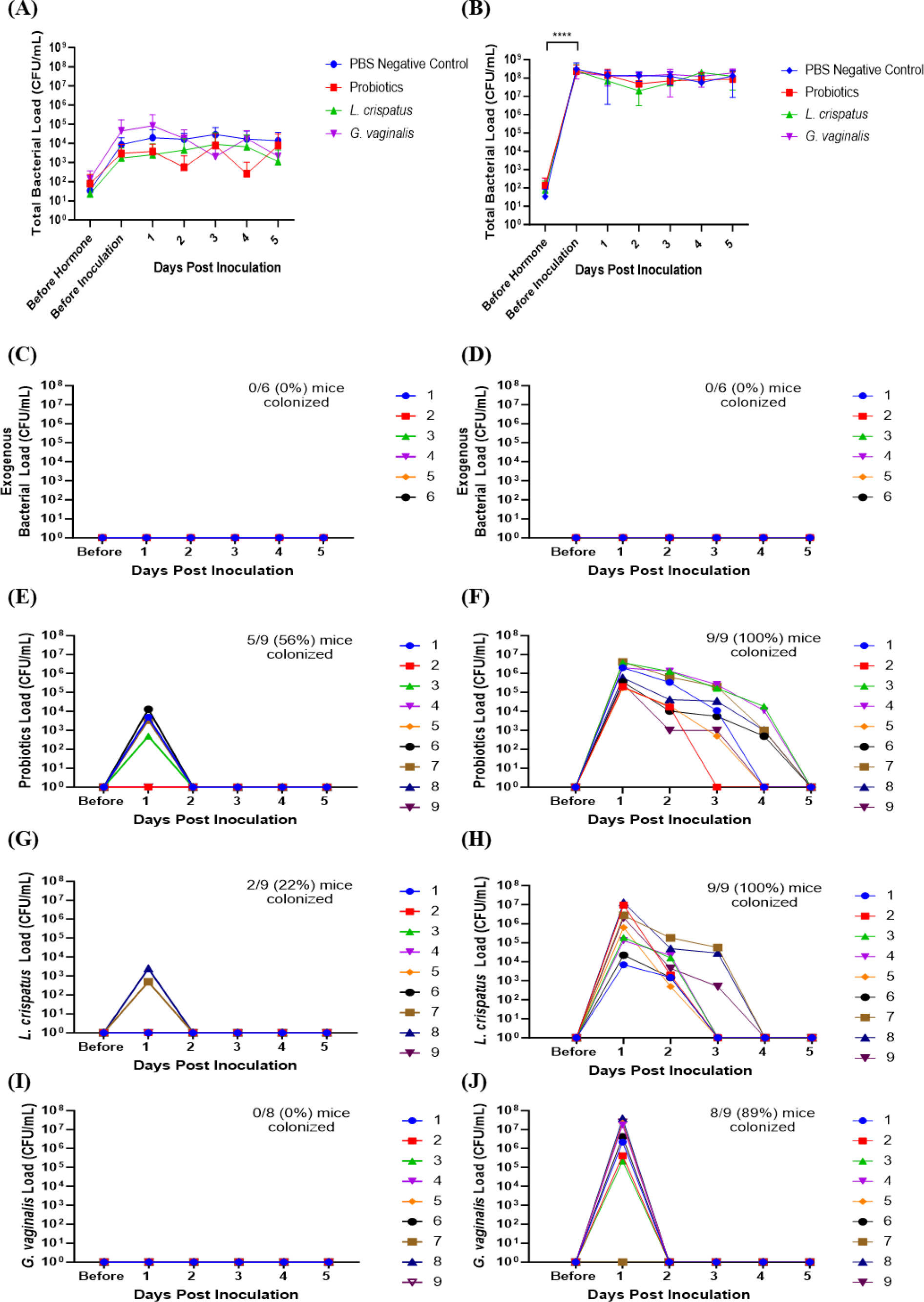
17β-estradiol treated OVX mice were fully colonized with human VMB species and had significantly higher bacterial loads compared to progesterone-treated OVX mice. Progesterone or 17β-estradiol 21-day release pellets were inserted subcutaneously into OVX mice 10 days after ovariectomies. Exogenous bacterial load and bacterial load were assessed by quantitative plating assays in mice before hormone treatment, 10 days after hormone treatment (before inoculation), and up to 5 days after one vaginal administration of a total of 10^7^ CFU *L. reuteri* RC-14 and *L. rhamnosus* GR-1, *L. crispatus, G. vaginalis*, or PBS as a no exogenous bacteria negative control. Data are n=9 per group, except PBS n=6, and are representative of two independent experiments. In the progesterone-treated mice, total colonies (A), exogenous bacteria colonies in PBS inoculated (C), probiotics colonies in probiotics inoculated (E), *L. crispatus* colonies in *L. crispatus* inoculated (G), and *G. vaginalis* colonies in *G. vaginalis* inoculated (I) were counted on agar plates. In the 17β-estradiol treated mice, total colonies (B), exogenous bacteria colonies in PBS inoculated (D), probiotics colonies in probiotics inoculated (F), *L. crispatus* colonies in *L. crispatus* inoculated (H), and *G. vaginalis* colonies in *G. vaginalis* inoculated (J) were counted on agar plates. In panels C, D, E, F, G, H, I and J, different mice are denoted by different colored points. The data was analyzed using two-way ANOVA with Tukey’s multiple comparisons (***p<0.001).

As expected, no mice in the PBS treated mice had any exogenous bacteria present (Figures 6C and 6D). Very few OVX mice treated with progesterone were colonized with any of the exogenous bacterial species administered for more than 24 h (Figures 6E, 6G, and 6I). Specifically, 5/9 (56%) of probiotics inoculated mice (Figure 6E), 2/9 (22%) of *L crispatus* inoculated mice (Figure 6G), and 0/9 (0%) of the *G. vaginalis* inoculated mice (Figure 6I) were colonized in progesterone-treated mice. However, 17β-estradiol-treated OVX mice were colonized with all the exogenous bacterial species (Figures 6F, 6H, 6J) for similar durations of time as normal (untreated) mice (Figure 1). Specifically, 9/9 (100%) of probiotics inoculated mice (Figure 6F), 9/9 (100%) of *L crispatus* inoculated mice (Figure 6H), and 8/9 (89%) of the *G. vaginalis* inoculated mice (Figure 6J) treated with 17β-estradiol were successfully colonized, similar to rates seen in normal mice (Figure 1). This indicates that the presence of 17β-estradiol is sufficient for colonization of human VMB in mice. As with normal mice (Figure 1), the total bacterial load did not exceed ∼10^7^-10^8^ CFU/mL after exogenous bacteria inoculation in the estradiol treated mice (Figure 6B), indicating the exogenous bacteria were displacing the endogenous bacteria and that there is a finite niche available in the vaginal environment. In progesterone treated mice, the available niche was much lower at ∼10^4^-10^5^ CFU/mL (Figure 6A).

### *Glycogen is upregulated in* 17β-estradiol *treated mice and MUC-1 is upregulated in progesterone treated mice*

Next, we examined a possible mechanism as to why 17β-estradiol promoted colonization with bacteria. We decided to look at two nutrients that are known to assist in colonization of vaginal bacterial. As previously mentioned, glycogen is a common nutrient source used by *Lactobacillus* species (Mirmonsef et al., 2014) and mucin is a common nutrient source used by *G*. vaginalis (Dupont, 2020; Vagios and Mitchell, 2021). Mice were ovariectomized, treated with 17β-estradiol or progesterone, and 10 days after hormone treatment, vaginal washes were collected and glycogen and Mucin-1 levels were measured. 17β-estradiol-treated mice had significantly increased levels of glycogen compared to progesterone-treated, OVX, and normal mice (Figure 7A) and progesterone-treated mice had significantly increased Mucin-1 levels compared to 17β-estradiol-treated mice (Figure 7B). Tissue samples were also collected ten days after hormone treatment and Mucin-1 staining was performed. Progesterone-treated mice showed significantly more mucin staining on the surface of the vaginal epithelium compared to other groups (Figure 7C), validating the results from the ELISA.

**Figure 7.**
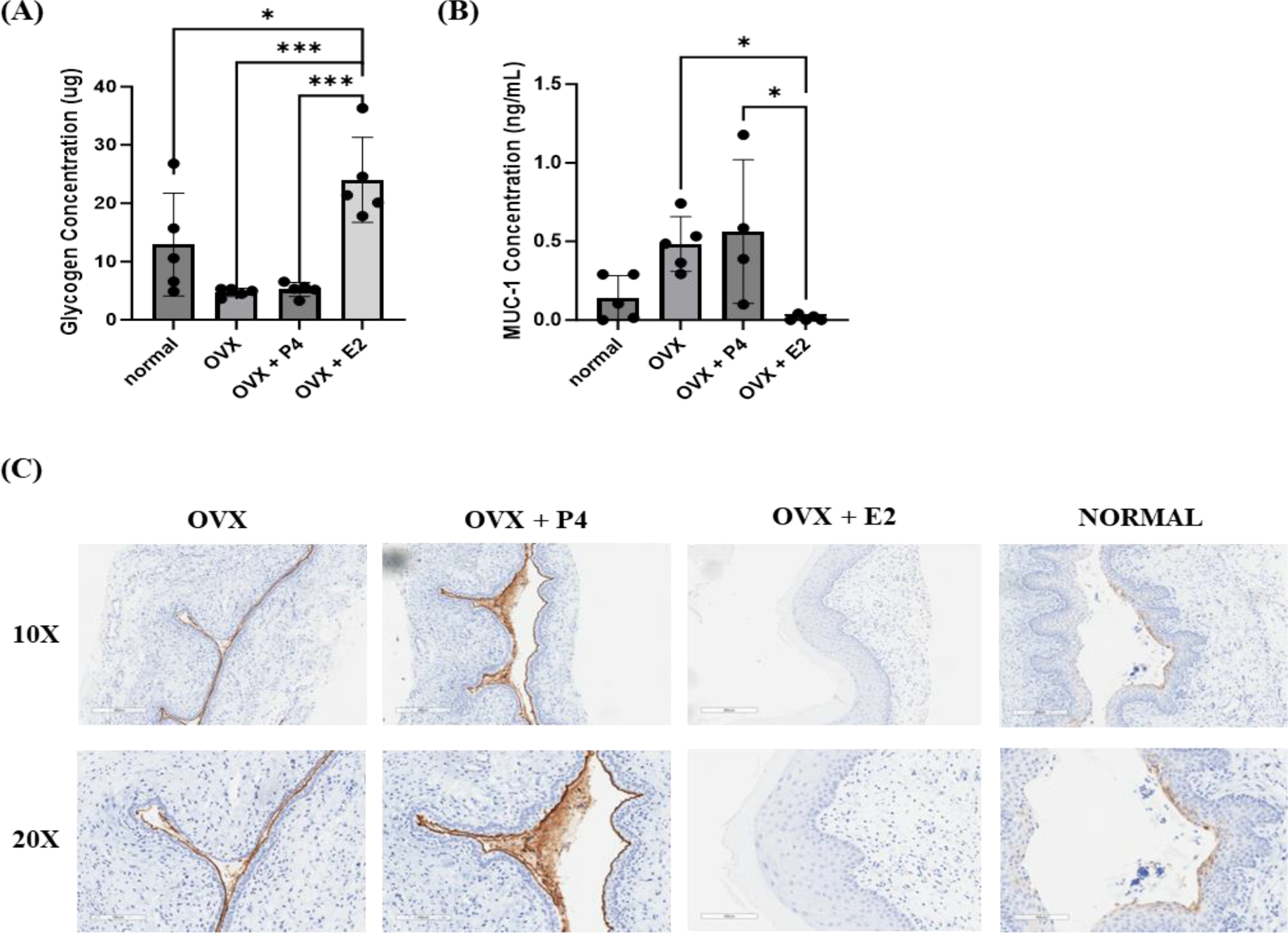
17β-estradiol-treated mice had increased glycogen levels and progesterone-treated mice had increased MUC-1 levels. 0.01 mg 17β-estradiol or 10 mg progesterone 21-day release pellets were inserted subcutaneously into female mice 10 days after ovariectomies. 10 days after hormone treatment, vaginal washes were collected from hormone-treated as well as OVX and normal mouse controls (n=5, representative of two independent experiments). Glycogen (A) and MUC-1 (B) levels were measured using a glycogen assay or a MUC-1 ELISA. The data was analyzed using one-way ANOVA with Tukey’s multiple comparisons (***p<0.001 and *p<0.05). 20 days after hormone treatment, mice were sacrificed, and vaginal tracts were collected from hormone-treated as well as OVX and normal mouse controls (n = 6, representative of two independent experiments). Vaginal tracts were fixed in methacarn for 72 hours and MUC-1 staining was performed. Positive staining for MUC-1 is brown.

## Discussion

In the past decade, there has been a great deal of interest in developing HMA mouse models that can be colonized with bacteria present in the human microbiota under optimal and dysbiotic conditions. These models can be used to study diseases and infections in the context of the microbiota composition and to understand the role microbiota health plays in disease outcomes (Arrieta et al., 2016). The majority of the studies conducted have been in the context of the gut microbiota. There are currently well-established animal models that mimic the human gut microbiota in mice (Hugenholtz and de Vos, 2018; Wrzosek et al., 2018). The establishment of these mouse models has been instrumental in advancing our understanding of the gut microbiota and the role it plays in various diseases such as inflammatory bowel disease (Gkouskou et al., 2014) and Alzheimer’s disease (Zhang et al., 2017). These models have also helped progress our understanding of the complex relationship between the gut microbiota and the immune system (Zheng et al., 2020). However, similar animal models of other human organ systems are greatly lacking. To study the female reproductive tract in the context of STIs, a similar mouse model that mimics the human VMB will be invaluable to study the mechanisms underlying VMB effect on STI susceptibility. Although a handful of studies have started to examine the establishment of such models (De Gregorio et al., 2015; Gilbert et al., 2019), this study is among the first to conduct an in-depth study in this area.

In this study, we successfully developed mouse models that mimic human eubiotic and dysbiotic VMB conditions. We first examined colonization in normal mice in their natural estrus cycle, to serve as a baseline. Other groups have also attempted to colonize hormone-unaltered mice with vaginal bacteria. One group attempted to colonize the mouse vaginal tract with human bacteria by inoculating vaginal swabs taken from women with BV into the vaginal canal of germ-free mice, however, the mice were not colonized with the bacterial species present in the swab from BV patients (Wolfarth et al., 2020). Another group developed a BV model in mice by colonizing mice with *Gardnerella vaginalis* and *Prevotella bivia*, two species commonly seen in BV patients. They were able to demonstrate that this coinfection model recapitulates several aspects of human BV, including vaginal sialidase activity, which is a diagnostic BV feature, epithelial exfoliation, and *P. bivia* ascending infection (Gilbert et al., 2013; Gilbert et al., 2019). Another study inoculated the mouse vaginal canal of BALB/c mice with different *Lactobacillus* species, including probiotic species *L. rhamnosus* and *L. reuteri*, and was able to successfully detect viable bacteria species up to four days post inoculation (De Gregorio et al., 2012). The limited success of these few studies demonstrates that while it is feasible to establish these models, many more studies need to be conducted to understand the conditions and factors for successful colonization before clinically relevant research questions can be addressed. Here, we described the conditions for successful, albeit temporary, colonization of normal mice. Furthermore, we identified the conditions for a dysbiotic model colonized with *G. vaginalis*, as well as a eubiotic model colonized with *L. crispatus* and a probiotic model colonized with *Lactobacillus* probiotic species *L. rhamnosus* GR-1 and *L. reuteri* RC-14. To our knowledge, this is the first study where the eubiotic CST I human VMB species *L. crispatus* was successfully detected in the mouse vaginal tract after inoculation.

Because we saw variations in the extent and duration of colonization in our model, we further examined known modifiable factors that could improve colonization. A number of studies have looked at the changes in the VMB of women throughout various stages of their life, including during puberty, menstruation, pregnancy, and menopause (Kaur, 2019). The shifts between these gynecological stages are largely regulated by fluctuations in sex hormones, indicating an underlying relationship between hormones and the changes in the VMB throughout a woman’s life. Therefore, we looked at the effect of hormones on bacteria colonization in our study and found many parallels with clinical data. We showed that removing endogenous hormones through OVX dramatically decreased VMB bacterial load compared to hormone-unaltered mice and neither eubiotic nor dysbiotic bacteria could colonize OVX mice for more than 24 h. Removing the effect of sex hormones through OVX may be considered congruent to menopause in women (Brzozowska and Lewiński, 2020). The transition of women through menopause has been shown to be marked by a gradual depletion of *Lactobacillus* species and an increase in anaerobic bacteria (Oliveira et al., 2022). Menopause is associated with a decrease in circulating hormones such as estrogen, which reduces glycogen deposition in the vaginal walls (Farage et al., 2010). Thus, it is likely that these changes lead to lower abundances of glycogen utilizing *Lactobacillus* in the vagina of postmenopausal women. Interestingly, estrogen hormone replacement therapy has been reported to increase *Lactobacillus* colonization in the VMB in post-menopausal women (Vitali et al., 2017; Geng et al., 2020), which is consistent with our findings. Treatment with 17β-estradiol restored the endogenous microbiota and colonization with eubiotic and dysbiotic bacteria in our mice. The results mentioned suggest the findings in our study recapitulate many aspects of the relationship between sex hormones and the VMB clinically, making it a reliable model to use in future studies.

The majority of the described effect of hormones on the VMB are from clinical studies. There are a few studies where sex hormones have been altered exogenously *in vivo* to assess the effect this has on the VMB. We have previously published showing humanized mice treated with MPA have increased VMB diversity (Wessels et al., 2019). As increased VMB diversity is associated with dysbiosis (Ravel et al., 2011; Juliana et al., 2020), this study indicates a link between progesterone and decreased VMB health. A few other groups have also published in this area. In one study, young female mice were administered 17β-estradiol or progesterone exogenously and vaginal bacterial loads were examined. They found that 17β-estradiol-treated mice had an increase in vaginal bacterial load, and progesterone treated mice had a near disappearance of all vaginal bacteria (Taylor-Robinson, 1991). As mentioned previously, in our study, we found similar results; mice in estrogen high states such as during estrus and 17β-estradiol-treated mice had increased colonization with exogenous bacteria compared to mice in progesterone high states. Another study administered 17β-estradiol and medroxyprogesterone acetate (MPA), a synthetic progestin, to mice and determined VMB contents (Priscilla Romina De Gregorio, 2018). They found the predominant taxa to be *Enterobacteria* in all experimental groups. They also found that lactic acid producing bacteria and *Enterobacteria* were found in greater concentration than *Staphylococci* and *Enterococci* in the 17β-estradiol treated groups. Finally, they found that higher numbers of cultivatable bacteria were present in 17β-estradiol treated mice than MPA treated mice, which is a similar finding to the previous study and our study as well. Another study administered probiotic *Lactobacillus* species to mice and found a greater number of viable bacteria during the proestrus–estrous stage of the estrus cycle compared to the metestrus–diestrus phase (De Gregorio et al., 2012), which is similar to our study as well. In the BV model Gilbert et al. developed with *G. vaginalis* and *P. bivia*, they treated mice with 17β-estradiol prior to colonization with BV-associated bacteria (Gilbert et al., 2019), however we were able to show colonization without this step, although estradiol completely restored colonization of all bacteria after OVX. Overall, these studies suggest that exogenous sex hormones may be altering the composition of the VMB, and that 17β-estradiol promotes overall bacterial load more than progestins *in vivo*. The results in our study align with previously published research in the area. To our knowledge, there are no comprehensive studies done where exogenous human eubiotic and dysbiotic bacterial species were administered to hormone-treated mice to show successful temporary colonization, which can be extended over time by repeated administration. To the best of our knowledge, this is the first study to optimize these conditions for prolonged colonization.

An additional optimization of our model was done by examining the effect of nutrient availability in the mouse vaginal canal, as this is an important determinant of what species colonize the VMB (Hood-Pishchany and Rakoff-Nahoum, 2021). As mentioned previously, glycogen is a common nutrient source used by *Lactobacillus* species (Mirmonsef et al., 2014) and mucin is a common nutrient source used by *G. vaginalis* (Dupont, 2020; Vagios and Mitchell, 2021). We found mucin to facilitate initial colonization by *G. vaginalis*. Studies have shown that women with BV have higher concentrations of mucin-degrading enzymes, which in turn decreases the vaginal fluid viscosity (Olmsted et al., 2003). Sialidase positive bacteria, such as *Gardnerella vaginalis*, are able to catabolize sialic acid in the cervicovaginal mucins and use it as a nutrient source, which could explain why the addition of mucin aided in *G. vaginalis* colonization in our model. In our study, mice treated with progesterone had increased Mucin-1 levels, which is a common nutrient source used by BV-associated bacteria, which aligns with published literature indicating a link between progesterone and increased diversity in the VMB (Wessels et al., 2019). Previous studies have also indicated progestins are associated with decreased glycogen production (Wessels et al., 2019), which is also congruent to our study. Whilst glycogen did not improve or hinder *Lactobacillus* colonization in our model, other studies have proposed that increased glycogen levels can decrease bacterial diversity and promote *Lactobacillus* colonization (Wessels et al., 2019). 17β-estradiol-treated mice did however have significantly increased levels of glycogen, which aligns with previously published studies (Gregoire and Parakkal, 1972; Mirmonsef et al., 2016). The results of our study suggest nutrient availability is largely dictated by sex hormones, which suggests a potential mechanism of how sex hormones affect the VMB.

In conclusion, this study provides new and important insights into the conditions that can facilitate colonization of the mouse vaginal tract with bacteria found in the human VMB. While clinical studies have provided significant insights regarding the correlation between colonization by different vaginal bacteria, mechanistic studies examining cause-effect relationships will require animal models, which are currently under-developed. Based on our results, estrogen appeared to play a critical role in creating a hospitable environment for colonization by all human vaginal bacteria, while substrates like mucin selectively enhanced colonization by specific anerobic species. The current model can enable further studies that directly examine the effect of the VMB on STI susceptibility such as HSV-2 and HIV-1, as well as examine the effect on other reproductive outcomes such as inflammation and epithelial barrier integrity. In the future, human microbiota associated mouse models will be invaluable tools to help further elucidate the complex role the VMB plays in vaginal health.

## Conflict of Interest

*The authors declare that the research was conducted in the absence of any commercial or financial relationships that could be construed as a potential conflict of interest*.

## Author Contributions

Conceptualization - CK and NR

Data curation – NR

Formal analysis – NR

Funding acquisition – CK

Investigation: NR, FM

Methodology – NR, FM, CK

Project administration – CK

Resources: CK

Software: NR

Supervision – CK, FM

Validation – CK

Visualization - NR

Writing – original draft – NR

Writing - review & editing – NR, CK

## Funding

This research was supported by grants from the Canadian Institutes of Health Research (CIHR Operating Grant FRN#126019 (C.K.); NR received funding support from the ontario graduate scholarship (OGS)

## Notes

### Competing Interest Statement

The authors have declared no competing interest.

